# Functional genomic characterization of anti-phage defense systems in vancomycin-resistant *Enterococcus faecium*

**DOI:** 10.64898/2026.11.03.764128

**Authors:** Eric Evans, Logan Tapia, Daria Van Tyne

**Affiliations:** Division of Infectious Diseases, University of Pittsburgh School of Medicine, Pittsburgh, Pennsylvania, USA; University of Pittsburgh School of Public Health, Pittsburgh, Pennsylvania, USA; Center for Evolutionary Biology and Medicine, University of Pittsburgh School of Medicine, Pittsburgh, Pennsylvania, USA

**Author notes:** Address correspondence to: Daria Van Tyne.

## Abstract

Vancomycin-resistant *Enterococcus faecium* (VREfm) is a significant public health threat, demonstrating high rates of antibiotic resistance and frequently causing hospital-acquired infections. Bacteriophages (phages) are a promising adjunctive therapy to reduce bacterial load and manage VREfm infections. Many bacteria encode anti-phage defense systems—groups of genes that can counteract phage infection—but these systems have not been characterized in VREfm. Here we explored the diversity of anti-phage defense systems within clinical VREfm isolate genomes and examined their impact on phage susceptibility. We first used two available bioinformatic tools (DefenseFinder and PADLOC) to identify putative anti-phage defense systems in 21 hybrid-assembled VREfm clinical isolate genomes, and observed that most anti-phage defense systems were located on plasmids, defense islands, and prophages. Combining this information with functional testing of phage susceptibility did not reveal robust associations between specific defense systems and resistance to four VREfm-targeting lytic phages. Due to inherent limitations of phage susceptibility testing, we next developed an assay to assess phage survival after co-incubation with bacteria, and identified over two dozen genes associated with phage killing, including a potentially novel defense island. Lastly, we attempted to functionally validate 10 candidate anti-phage defense systems, only four of which were ultimately tested in enterococci. Expression of one of these candidates, AbiE, decreased bacterial susceptibility to two *Enterococcus*-targeting phages. As VREfm- targeting phage therapy becomes more widely available, identification of genes contributing to phage resistance can aid in streamlining the provision of phages to patients who could benefit from this new therapeutic modality.

**IMPORTANCE:** VREfm is a significant public health threat, acquiring resistance to last-line antibiotics and becoming endemic in healthcare systems. There is a clear need for alternative therapies to treat VREfm infections. In this study we identified and characterized genes that are predicted to cause resistance to bacteriophages, an adjunctive therapeutic approach for treating antibiotic-resistant bacterial infections, among VREfm collected from infected patients. Our findings reveal a wide diversity of anti-phage defense systems in clinical VREfm isolates, and identify several possible drivers of phage resistance in this organism. The knowledge gained will inform the development of phage therapy for patients with difficult-to-treat VREfm infections.

## INTRODUCTION

*Enterococcus faecium*, a common nosocomial pathogen, has evolved from a commensal bacterium to become both multidrug-resistant and endemic in many healthcare systems (1–3). Causing over 54,000 hospitalizations and 5,400 deaths in 2017, vancomycin-resistant *E. faecium* (VREfm) is a leading cause of healthcare-associated infections (4). Mortality rates have remained consistently high, with the 30-day mortality rate of VREfm bloodstream infections hovering around 30% for the last 25 years (3, 5–9). VREfm also disproportionately affects immunocompromised and hospitalized individuals and is a significant public health threat (10). Altogether, there is a clear need for the develop new treatments for VREfm infections.

Bacteriophages (phages) are viruses that infect bacteria, using the host cellular machinery to replicate before lysing the cell (11). The use of phages to treat bacterial infections has recently gained traction as an alternative or adjunctive to conventional antibiotics. Phage therapy has been used to reduce bacterial load and improve bacterial clearance, including in enterococcal infections (12–14). Genus- or species-specific phage libraries are often needed for phage therapy (15, 16), because phages often have narrow host ranges (17–19). Despite the high potential of phage therapy as a new treatment option, narrow specificity can be an obstacle to the rapid provision of phages for widespread treatment.

There are several factors that determine whether a given bacterial cell is amenable to productive phage infection. Phages are often dependent on bacterial surface decorations to bind to and infect a host cell, effectively being limited by the availability of cell surface receptors for phage infection (17, 20). Additionally, bacteria often contain anti- phage defense systems, a type of immune response to prevent phage replication (21, 22). Anti-phage defense systems can cleave foreign DNA, reduce available energy, block protein production, and even kill the infected bacteria through toxin activation (21, 22). The roles of many anti-phage defense systems in phage resistance have yet to be fully characterized.

In this study, we assessed the diversity and functional impacts of putative anti- phage defense systems among contemporary VREfm clinical isolates. We determined the susceptibility of 21 VREfm clinical isolates to four *Enterococcus*-targeting phages, used a new approach to assess phage viability upon infection of clinical isolates, and identified both previously discovered and novel putative anti-phage defense systems associated with phage-mediated killing of VREfm.

## RESULTS

### Anti-phage defense systems in VREfm clinical isolates are diverse and often encoded on mobile elements

We performed short- and long-read whole genome sequencing and hybrid genome assembly of 21 VREfm clinical isolates from bloodstream infections collected from patients at UPMC (**Supplementary Table 1**). We then used DefenseFinder and PADLOC to search for putative anti-phage defense systems in all 21 hybrid assembled genomes. We observed moderate agreement between these search tools, with approximately half of all defense systems identified by both tools, 24.4% of systems identified uniquely by DefenseFinder, and 25.4% of systems identified uniquely by PADLOC. Each genome encoded 4-13 predicted defense systems. The AbiE abortive infection system was the most common system identified, with all 21 (100%) isolates having at least one copy (**Figure 1A**). Other prevalent systems included mazEF, which were observed in 20 (95.2%) isolates, and PD-Lambda-1, which was found in 18 (86.7%) genomes. We next performed manual curation to assess each anti-phage defense system and ensure that the identified genes appeared to be complete and intact in the genome. This manual curation revealed that several systems were interrupted in one or more genomes, including Abi2, BREX, Gabija, RM, and PD-T7-3 (**Figure 1A**). Overall, we observed that 7.0% of all defense systems identified were interrupted and likely non-functional, suggesting that defense system identification tools do not always identify intact systems. Nonetheless, these data demonstrate a wide array of defense systems among VREfm clinical isolate genomes.

**Figure 1.**
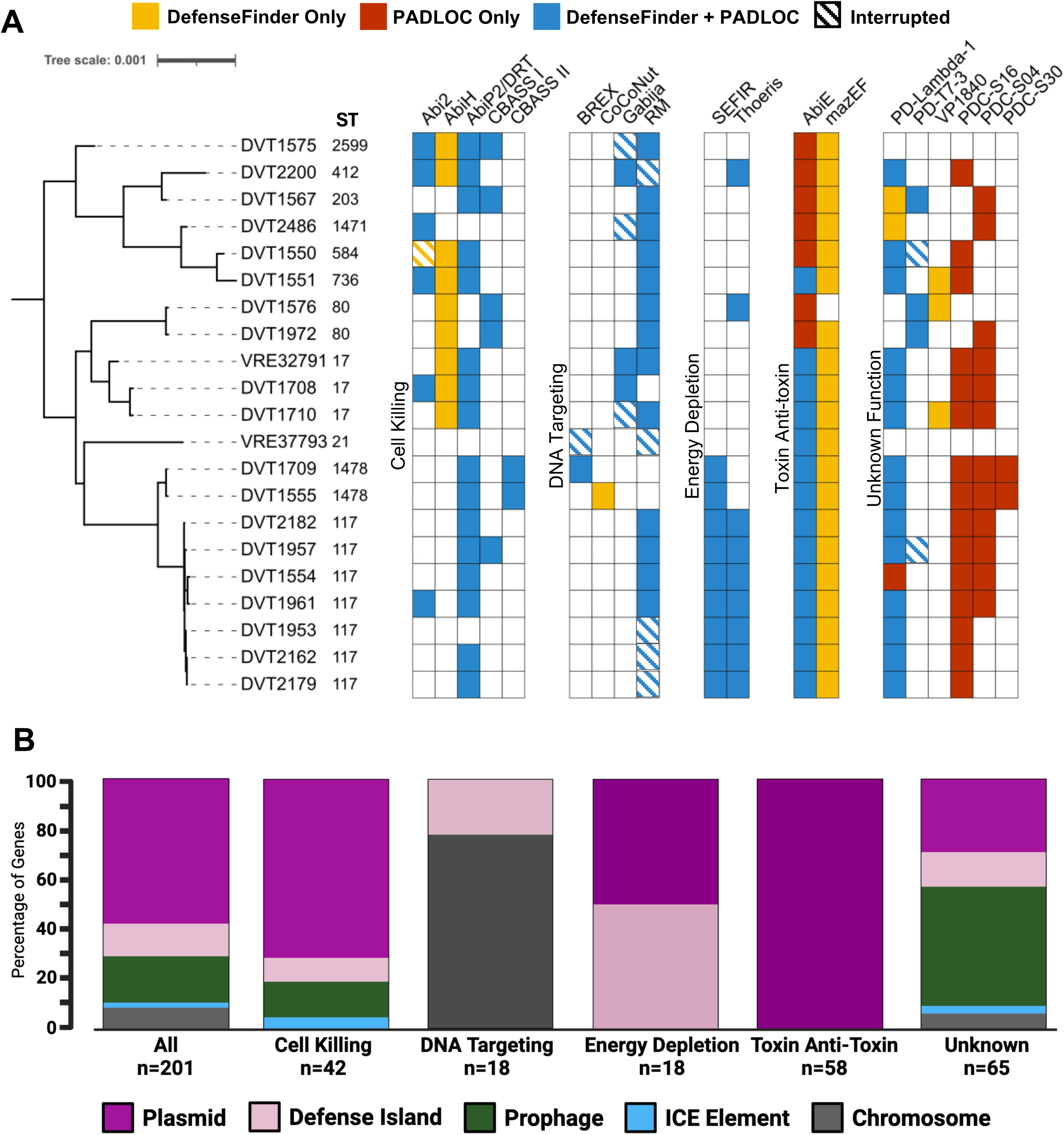
Anti-phage defense systems identified in VREfm clinical isolates and their genomic locations. (A) Anti-phage defense systems found in 21 representative clinical VREfm isolate genomes. Isolates are organized by their placement on a core genome phylogenetic tree. Defense systems were identified with DefenseFinder and PADLOC, followed by manual curation to identify systems which were interrupted and predicted to be non-functional. ST = sequence type. (B) Graphical visualization of the genomic location of intact anti-phage defense systems, organized by defense system type and plotted as relative proportion of systems found on different types of mobile elements, or not mobile element-associated (Chromosome).

After the locations of putative anti-phage defense systems were recorded, the bacterial genomes were assessed to identify mobile genetic elements encoding anti- phage defense systems through co-localization. We characterized the location of each intact defense system as being on the chromosome (i.e., not mobile element-associated) or on a mobile genetic element, including plasmids, defense islands (i.e., mobile elements enriched in defense systems, detailed in **Supplementary Figure 1**), prophages, and integrative conjugative elements (ICEs). We observed that over 90% of the 201 intact anti-phage defense systems were located on mobile genetic elements, with 57.7% (n=116) found on plasmids, 12.9% (n=26) found on defense islands, 18.4% (n=37) found on prophages, and 2.0% (n=4) found on ICEs (**Figure 1B**). The genomic locations of anti- phage defense systems were different between different categories of defense systems. For systems related to cell killing, the vast majority (71.4%) were found on plasmids, while systems targeting foreign DNA were primarily not mobile element-associated (77.8%).

Anti-phage defense systems targeting phages through energy depletion were evenly split between plasmids and defense islands, and toxin-anti-toxin systems were always found on plasmids, consistent with their origin and common functions (23). Of the remaining anti-phage defense systems of unknown function, nearly half (47.7%) were found on prophages. Taken together, these data demonstrate that while the vast majority of anti- phage defense systems are mobile element-encoded, their precise genomic locations vary by system type.

### Susceptibility to four lytic phages is not correlated with anti-phage defense system presence

To investigate associations between putative anti-phage defense systems and phage susceptibility or resistance, we screened all 21 VREfm isolates for susceptibility to four *Enterococcus*-targeting phages, called Hi3, 113, Ben, and Bop (**Figure 2A, Supplemental Table 1**). Hi3 had the broadest host range, showing plaquing activity on 71.4% (n=15) of isolates. 113 was the least active phage, plaquing on only 19.1% (n=4) of isolates. The other two phages, Ben and Bop, demonstrated similar patterns of activity, with both phages showing lytic activity on over half of the isolates. We compared the total number of anti-phage defense systems among isolates that were susceptible or resistant to each phage, and did not observe any striking differences (**Supplementary Figure 2**). Using the spot titering data, we next tested for associations between putative anti-phage defense system presence and phage resistance using Fisher’s Exact Test to compare the proportion of isolates with a specific anti-phage defense system that were resistant with the proportion of isolates that were susceptible or showed lytic activity only. The strongest association detected was between Gabija and resistance to phage Ben (P=0.0143, **Figure 2B**), though this association was considered weak because the P- value was not significant after correcting for multiple comparisons. The lack of statistically significant results in this analysis could be due to the small sample size, functional redundancy between anti-phage defense systems, or confounding from bacterial population structure.

**Figure 2.**
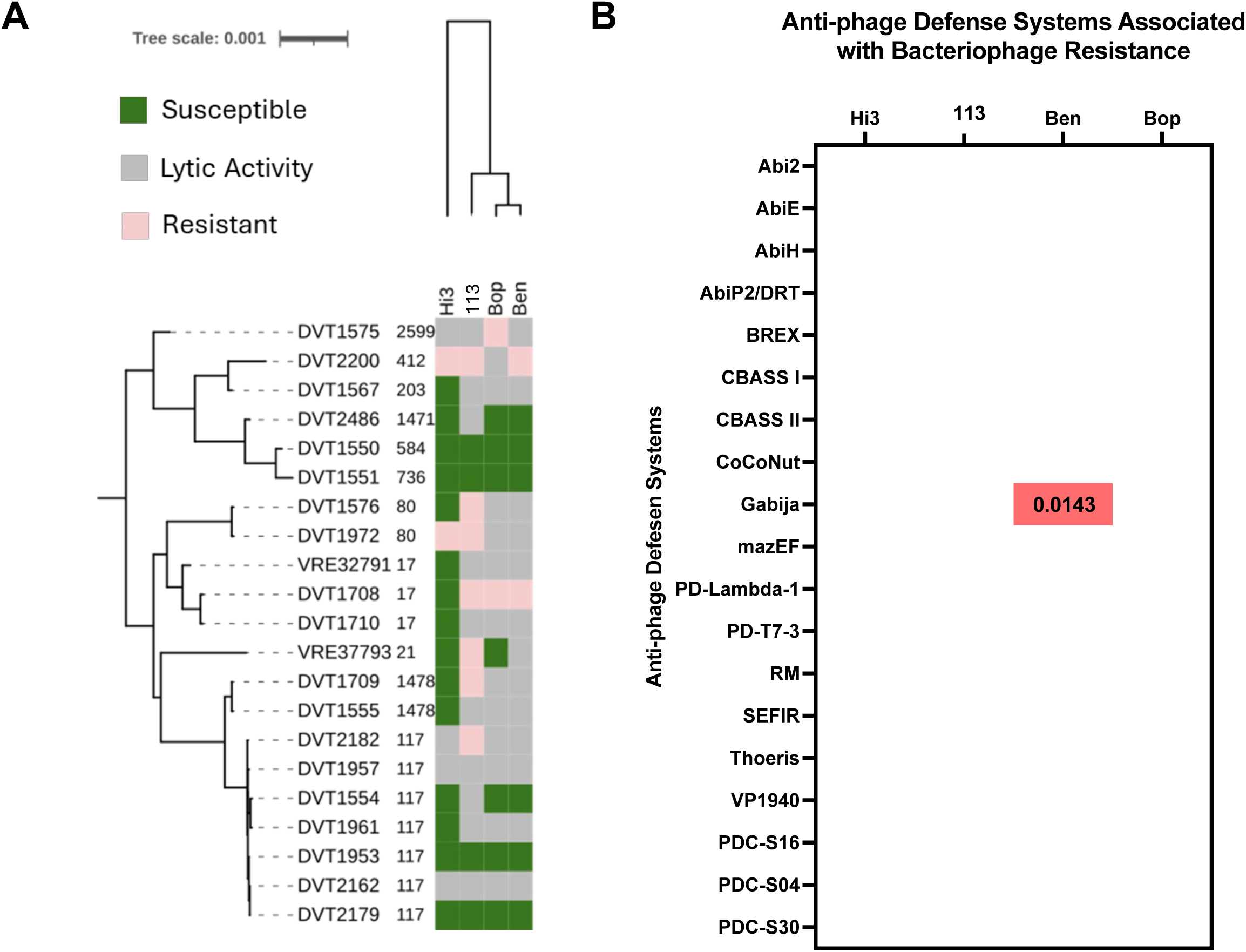
VREfm phage susceptibility and associations with anti-phage defense systems. (A) Susceptibility of 21 VREfm clinical isolates to four *E. faecium*-targeting lytic phages. Susceptible = phages formed countable plaques; Lytic Activity = bacteria lysis but no plaques observed. Resistant = no detectable phage activity. (B) P-values for associations between anti-phage defense system presence and resistance to each of four tested phages. Fisher’s Exact test was run for every defense system-phage pairing, and naïve P-values <0.05 are shown.

### Measuring phage killing and survival identifies known and novel genes associated with phage killing

Phage susceptibility testing revealed that for many isolates, the tested phages demonstrated lytic activity without clear plaquing, suggesting that the phages were killing the bacteria but were unable to replicate and form plaques, perhaps because they trigger abortive infection or other anti-phage defense mechanisms. To better characterize this activity, we quantified the total amount of phage present before and after four hours of incubation with each bacterial isolate, calling this a “phage survival” assay. By monitoring the abundance of active phage rather than bacterial survival, this assay enabled us to assess whether each phage was able to infect and replicate (as evidenced by an increase in phage abundance after four hours), was unable to infect (no change in phage abundance), or was targeted by the bacteria for inactivation or killing (decrease in phage titer). When we compared the data from this assay with the previous spot titering data, we observed that for the highly active phage Hi3, most phage-susceptible phage-isolate pairings showed a greater than 1-log_10_ increase in phage abundance over the four-hour period, suggesting productive phage replication (**Figure 3A**). None of the pairings in the “lytic activity” or “resistant” categories showed a decrease in phage abundance that would suggest killing of Hi3. When isolates were exposed to phages 113, Ben, or Bop, however, several of the “lytic activity” and “resistant” phage-isolate pairings demonstrated greater than 1-log_10_ decreases in phage titer, suggesting inactivation or killing of the phages. For these three phages, susceptible pairings also demonstrated increases in phage abundance compared to the other categories (**Figure 3A**). Taken together, these results suggest that conclusions about phage susceptibility or resistance likely depend on the assay used.

**Figure 3.**
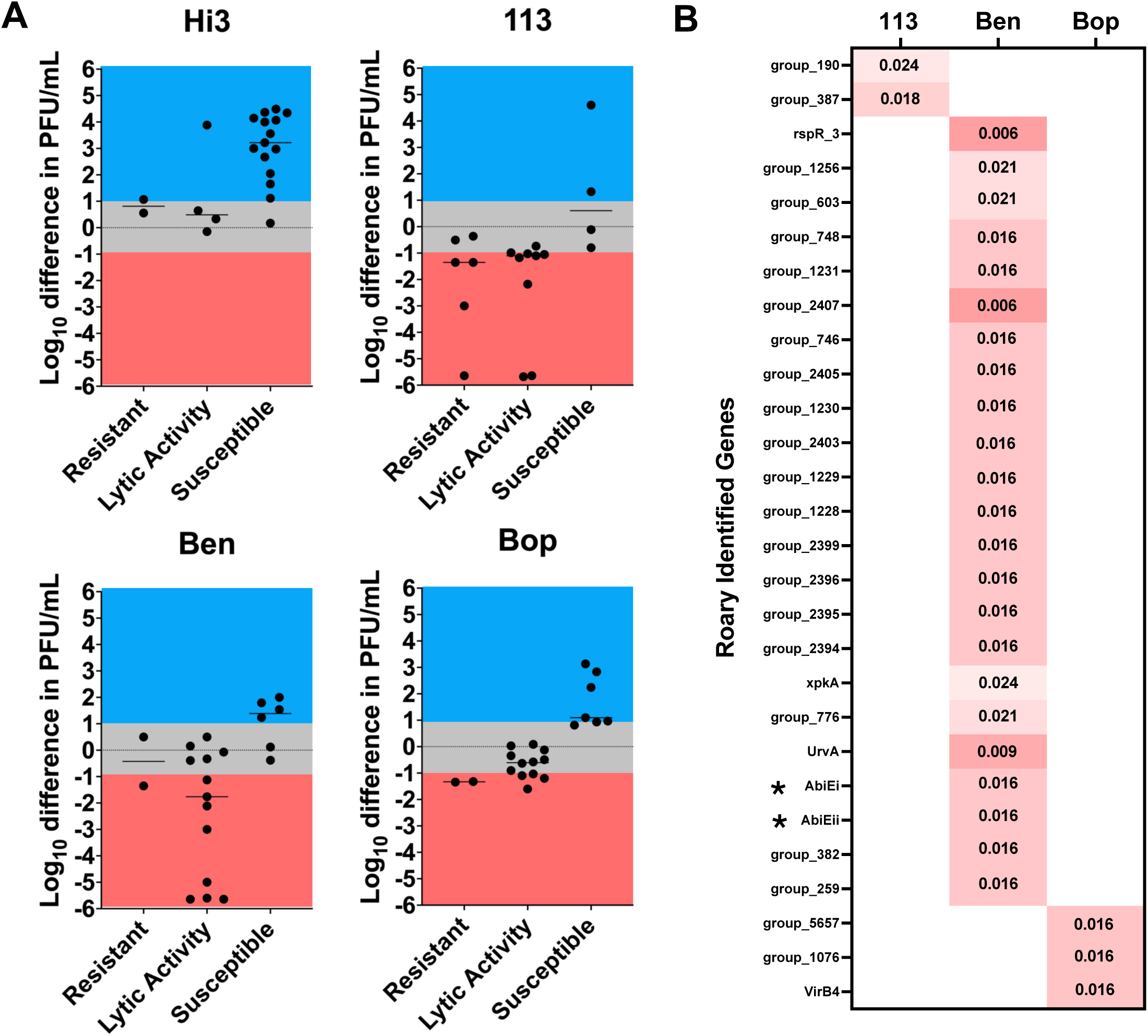
Phage survival assay identifies known and novel putative anti-phage defense systems. (A) Comparison of phage survival assay data (shows as log_10_ differences in PFU/mL between T=0 and T=4-hour timepoints) with phage susceptibility data as measured in a standard titer assay. Blue indicates phage titer increase due to productive infection, while red indicates phage titer decrease and possible phage killing. Two isolates were removed from the phage 113 dataset due to inconclusive results. (B) P-values for associations between anti-phage defense system presence and phage killing for phages 113, Ben, and Bop. Naïve P-values <0.025 from scoary analysis are shown. Asterisks indicate putative anti-phage defense system genes.

Using the data from the phage survival assay, we first looked for associations between the presence of individual anti-phage defense systems and phage killing, but did not identify any significant associations. Next, we took an unbiased approach to identify genes in the bacterial genomes that were associated with phage killing using scoary. As no isolates demonstrated a decrease in phage abundance upon incubation with Hi3, this analysis was only performed for phages 113, Ben, and Bop. A total of 28 genes were found to be associated with phage killing (naïve P<0.025) (**Figure 3B, Supplementary Table 2**). While a large proportion of associated genes were identified as encoding hypothetical proteins, two different alleles of the previously identified anti-phage defense system AbiE (AbiEi, AbiEii) were found to be associated with killing of phage Ben (naïve P=0.016) (**Figure 3B**). Also of note, 13 genes associated with killing of phage Ben were found to co-localize within a gene cluster reminiscent of a defense island (**Supplementary Figure 3, Supplementary Table 2**). Taken together, these data suggest that currently identifiable anti-phage defense systems are not robustly associated with protection from the tested phages, and that there could be other genes that contribute to phage resistance and phage killing in VREfm that have not yet been explored.

### Functional validation of anti-phage defense systems in *Enterococcus* shows AbiE contributes to phage resistance

Having explored phage resistance and its relationship to anti-phage defense systems, we next turned to functional validation of anti-phage defense system activity. We selected 6 anti-phage defense systems as candidates for molecular cloning, four of which had two different alleles among clinical VREfm isolates (**Supplementary Table 3**). We first attempted to clone each of the 10 anti-phage defense systems into *E. coli,* followed by transformation into the *E. faecium* strain ATCC 19434, however these attempts were largely unsuccessful (**Figure 4A**). Only four of the ten constructs were able to be cloned into *E. coli*, and of these only one was successfully transformed into *E. faecium.* We then selected the remaining nine anti-phage defense systems to attempt cloning and functional validation directly in the *E. faecalis* OG1RF strain, which resulted in successful cloning of three additional systems (**Figure 4A).** Of the four systems successfully expressed in enterococci, three (CBASS I, Gabija, and AbiP2 Type 1) showed no change in phage susceptibility for any of the tested phages. OG1RF transformed with AbiE Type 2, however, showed a visibly reduced titer and a switch to lytic activity against phages Ben and Bop, suggesting that expression of this defense system might have triggered abortive infection upon phage exposure (**Figure 4B**). While the AbiE allele tested was different from the alleles associated with phage killing by scoary, these data nonetheless suggest that AbiE plays a role in defending from phage infection.

**Figure 4.**
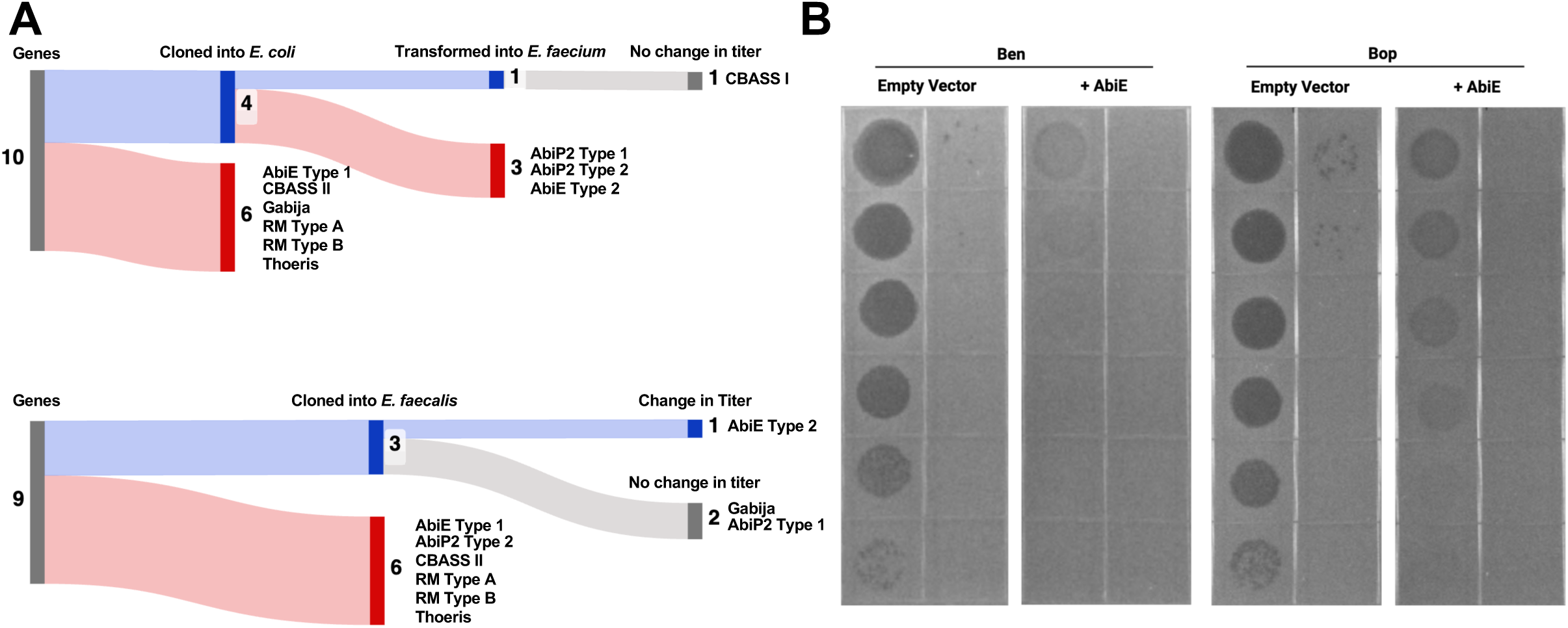
Functional validation of predicted anti-phage defense systems in *Enterococcus*. (A) Schematic diagrams of attempted functional validation of 10 putative anti-phage defense systems. Cloning in *E. coli* followed by transformation into *E. faecium* is shown in the top diagram, while direct cloning into *E. faecalis* is shown in the bottom diagram. Blue shading indicates successful attempts while red shading indicates failed attempts. Grey shading indicates when functional testing showed no change in phage titer. (B) Soft agar overlay spot titering of phages Ben and Bop against *E. faecalis* OG1RF carrying empty vector or AbiE Type 2.

## DISCUSSION

The aim of this study was to characterize the diversity of anti-phage defense systems in clinical VREfm isolates and to identify systems that might drive phage resistance in this pathogen. Using multiple approaches, we searched for correlations between anti-phage defense system presence and resistance to phage-mediated killing. While no strong correlations between specific anti-phage defense systems and phage resistance were found, several known and novel putative defense genes were found to be weakly associated with phage resistance and/or killing. Functional validation of 10 candidate anti-phage defense systems yielded only one confirmed hit, suggesting that these systems are either highly specific to the phage(s) against which they can protect, are dependent on factors that were not tested here, or are not in fact involved in phage defense in VREfm.

We used two different but widely cited bioinformatics tools to identify putative anti- phage defense systems in each VREfm genome. We were surprised that only about half of all systems were identified by both tools, but this could be due to the different databases and search parameters used. Regardless of the search tool used, subsequent manual curation removed nearly 10% of the identified genes because they were interrupted by insertion sequences or contained other variants likely to inactivate them. This suggests that current methods for defense system identification do not always properly assess the completeness of these systems, and perhaps should be refined.

We were also surprised to observe a lack of strong associations between anti- phage defense system presence and phage susceptibility, as measured by conventional soft agar overlay spot titering. For this reason, we decided to focus instead on quantifying phage abundance after incubation with each bacterial isolate to distinguish potential contributors to the “killing” of each phage. This phage survival data was helpful in revealing instances of phage killing among phage-isolate pairings demonstrating both lytic activity and phage resistance in the soft agar overlay assay. Phage killing data were generally correlated with spot titering data, with no susceptible pairings showing evidence of phage killing. When we used the phage survival assay data to search for other potential genetic contributors to phage killing beyond currently identifiable anti-phage defense systems, we identified two AbiE alleles and a putative novel defense island that were associated with killing of phage Ben. A recent study also used phage outcome (i.e., burst size) to assess anti-phage activity, suggesting that this could be a worthwhile approach (24). The AbiE alleles were identified as present by DefenseFinder and PADLOC, but were not significantly associated with phage killing in our initial analysis, likely because that analysis grouped all AbiE alleles together. We also identified two genes associated with phage 113 killing, three genes associated with phage Bop killing, and 21 genes associated with phage Ben killing. Exploring the role of these genes in protection from phage-mediated killing will be a focus of our future work.

Attempts to express individual anti-phage defense systems in *E. faecium* proved to be largely unsuccessful. Most of these cloning efforts failed while attempting to clone plasmids in *E. coli*, perhaps because aberrant defense gene expression caused bacterial cell death. Removing the inducible expression system of pMSP3535 and transforming putative defense systems directly into enterococci allowed us to functionally test more candidate systems. However, the anti-phage defense systems we were able to successfully transform into enterococci displayed varied effects on phage killing. While transformants containing CBASS I, AbiP2/DRT, and Gabija did not exhibit changes in phage susceptibility, the transformant expressing AbiE Type 2 did show a change, demonstrating reduced phage titer and a change to lytic activity only for both phages tested, similar to other reports (21, 25–27). While functional validation was achieved for a single AbiE allele, this was only one of the four systems tested and one of 10 genes that we attempted to validate, suggesting that only a small percentage of putative defense systems are verifiable with this approach.

This study had several limitations. First, our sample size of both VREfm clinical isolates and *Enterococcus*-targeting phages were both rather small. The isolates we included nonetheless spanned the current diversity of VREfm clinical isolates, and the small number of phages tested reflects the fact that there are not currently large numbers of lytic phages available that target VREfm. Additionally, we only attempted to functionally validate a relatively small number of the defense systems identified, as molecular cloning and transformation both proved arduous tasks. Nonetheless, our findings suggest that predicted anti-phage defense systems are just that; predictions that should be studied further.

Bacteriophage therapy is a potentially powerful adjunctive therapy in the fight against antibiotic-resistant bacterial infections. While most of the available literature to date investigating anti-phage defense systems has focused on *E. coli*, this study advances the field by exploring and functionally validating anti-phage defense systems inVREfm, a prominent human pathogen. Further exploration of these systems can inform the treatment of patients with VREfm-targeting phages, and could uncover new molecular tools for the study of VREfm in clinical settings.

## MATERIALS AND METHODS

### Bacterial Isolate and Phage Collections

Bacterial isolates used in this study were collected from the University of Pittsburgh Medical Center (UPMC) system. Once collected from the clinical microbiology laboratory, isolates were grown in BHI overnight and stored in 16% glycerol at −80°C. (**Supplementary Table 1**) (28). Collection of clinical isolates from UPMC was approved by the Institutional Review Board at the University of Pittsburgh under protocol STUDY21040126, with some isolates being described previously (29) (**Supplementary Table 1**). Phages used in the study were originally isolated from wastewater. *Enterococcus* phage 113 (ATCC 19950-B1) was purchased from ATCC (30), Hi3 was kindly provided by the Duerkop lab (14), and VB_OCPT_Ben (Ben) and vB_OCPT_Bop (Bop) were kindly provided by the Whiteson lab (31).

### Genome Sequencing and Analysis

For phage genomic DNA extraction, 250µL of a high-titer phage lysate was mixed with 0.25µL DNase (1000U/mL) and 0.25µL RNase (10mg/mL) and incubated at 37°C for 10 minutes. A Qiagen DNeasy Blood and Tissue kit (Qiagen) was then used following the manufacturer’s Quick Start Protocol. For bacterial genomic DNA extraction, bacterial isolates were grown overnight in Brain Heart Infusion (BHI) broth at 37°C, then 1 mL of overnight culture was centrifuged at 13,000 RPM for 2 minutes and the pellet was resuspended with 1mL of TE buffer (10 mM Tris-HCl and 1 mM EDTA, pH 8.0). This step was repeated twice. Lysozyme (50µL at 50mg/mL) and mutanolysin (10µL at 2500U/mL) were added, and the mixture was vortexed and incubated at 37°C for 1 hour, and then a Qiagen DNeasy Blood and Tissue kit was used following the manufacturer’s Quick Start Protocol. DNA concentrations were measured using an Invitrogen Qubit 4 Fluorometer (ThermoFisher Scientific) before library construction and sequencing on the Illumina platform (2 x 150 bp paired-end reads, sequencing performed by SeqCenter, LLC). Long- read sequencing was performed on extracted gDNA using an Oxford Nanopore Technologies Mk1C device with R10.4.1 (FLO-MIN114) flow cells and SQK-RBK114.24 rapid gDNA barcoding kit. Basecalling and demultiplexing were performed with Dorado v0.9.6 (high accuracy model), specifically dna_r10.4.1_e8.2_400bps_hac@v5.0.0. Hybrid assembly was performed with Unicycler v0.5.1 (32).

Phage genomes were assembled using SPAdes v3.15.5 with the --metaviral flag (33). Phage genome dendrograms were generated with VIPTree v4.0 (34), and were visualized using iTOL v7.2 (35). Bacterial MLST was assessed using mlst v2.11 (36). Bacterial whole genome phylogenetic trees were generated by annotating each genome with prokka v1.14.5 (37), orthoclustering and core genome alignment with roary v3.13.0 (38), and tree construction with RAxML v8.2.12 using the GTRGAMMA algorithm and 100 replicates (39). Whole genome sequencing data for phages and bacterial strains are deposited at the National Center for Biotechnology Information (NCBI), with accession numbers listed in **Table 1** and **Supplementary Table 1**.

### Phage Susceptibility Testing

Phage activity was assessed using the soft agar overlay spot titering method on Todd Hewitt agar plates (40). Briefly, serial dilutions of each phage in SM+ buffer (50mM Tris HCl, 100mM NaCl, 8mM MgS04, and 5mM CaCl2) were spotted onto a soft agar overlay of each bacterial isolate, grown overnight at 37°C, and visualized the following day for the appearance of lytic plaques. Isolates that showed individual plaques at any dilution were designated as phage-susceptible. Isolates showing zones of clearance but having no observable plaques were designated as having phage “lytic activity”. Isolates without any clearance of the bacterial lawn were designated as phage-resistant.

### Anti-phage defense system Identification and Visualization

The DefenseFinder v2.0.1 tool was run with coverage set to 0.5 to identify putative anti-phage defense systems present within each bacterial isolate genome (41–44). Identification of anti-phage defense systems was also performed using PADLOC v2.0.0 (45, 46). Defense system genome coordinates were used to assess completeness of anti- phage defense system sequences and locate associated mobile genetic elements using Geneious Prime v2025.0.3. Visualization of loci surrounding anti-phage defense systems was performed using EasyFig v2.2.5 (47).

### Identification of Mobile Genetic Elements Encoding Anti-phage Defense Systems

Plasmids encoding anti-phage defense systems were identified using visualization of each bacterial genome in Geneious Prime v2025.0.3. Conserved regions found on the main chromosome within at least one isolate that contained more than 10 genes, had a putative anti-phage defense system present, and had at least five genes flanking either side that were consecutively conserved in other isolates were considered to be defense islands (21, 22). Defense islands were identified using the gene_presence_absence output file from roary v3.13.0 (48). Locations of anti-phage defense systems were marked, and the flanking genes were analyzed for conservation across multiple VREfm isolates. Prophages were identified with PHASTEST v3.0 (49). Integrative conjugative elements (ICEs) were identified using ICEberg v3.0 (50).

### Cloning of Anti-phage defense systems

Individual anti-phage defense systems were amplified and cloned into the pMSP3535 inducible expression vector, or a pMSP3535-derived shuttle vector, using Gibson assembly (**Supplementary Tables 3 and 4**) (51). The pMSP3535-derived shuttle vector was created by removing the nisin-dependent promoter region through amplification of the remaining vector and reassembly using Gibson assembly. Genes cloned into the shuttle vector were cloned with 150bp of the upstream sequence to capture native promoter elements. Information on vector and anti-phage defense system combinations can be found in **Supplementary Table 4**. Gibson assembly products were transformed into NEB 5-alpha *E. coli* cells and plated onto Luria Broth (LB) agar with 300 ug/mL erythromycin. Individual colonies were selected for colony PCR using GoTaq Green with “check” primers listed in **Supplementary Table 4**. Positive colonies were grown overnight in LB with 300 µg/mL erythromycin to amplify the vector prior to extraction with a HiSpeed Plasmid Maxi Kit (Qiagen) following the manufacturer’s protocol. Plasmid DNA concentrations were measured using a NanoDrop ND-1000 Spectrophotometer (Thermo Scientific). Approximately 1 µg of the amplified plasmid was then transformed into ATCC 19434, an *E. faecium* laboratory strain, or OG1RF, an *Enterococcus faecalis* laboratory strain, which were made to be electrocompetent (52). Transformed bacteria were plated on Brain Heart Infusion (BHI) agar containing 15 µg/mL erythromycin. After growth of colonies was observed, colonies were checked by colony PCR with GoTaq Green to confirm the vector with the correct insert was present using check primers listed in **Supplementary Table 4**. Positive colonies were then stored in BHI media with 16% glycerol at −80°C for further experimentation.

### Phage Survival Assay

Bacteria from overnight cultures (600 µL) were added to 12 mL of Todd Hewitt Broth and allowed to grow until an OD600 of approximately 0.3. 3 mL aliquots were removed, and individual phages were added to separate aliquots at an MOI of 0.01. At timepoint 0, 1 mL was immediately taken and filtered through a 0.22 µm filter and stored at 4°C. Bacteria and phage were allowed to grow together for 4 hours, then 1 mL was removed, filtered, and stored at 4°C. Filtrates were serially diluted in SM+ buffer, spot titered onto the host of each phage, incubated overnight, and then phage plaques were counted to determine the plaque-forming units (PFU) per mL at each time point.

### Identification of Bacterial Genes Associated with Phage Killing

Gene_presence_absence outputs from roary were used to perform a scoary v3.13.0 analysis of bacterial genes associated phage killing for each of the phages used (53). For this analysis, roary was run with the standard minimum percentage identity of 0.95. Naïve P-values generated by scoary (p<0.025) were used to identify genes associated with phage killing (**Supplementary Table 2**).

### Statistical Analyses

Differences in the number of anti-phage defense systems found between different groups of isolates were assessed using the Mann-Whitney U test. Differences in phage susceptibility between groups of isolates that did or did not encode a given anti-phage defense system were analyzed using Fisher’s Exact Test, with a P-value lower than 0.05 noted as suggesting an association. Associations between accessory genes and phage killing were assessed with scoary v3.13.0, with associations with P-values less than 0.025 reported.

## Data Availability

WGS data are available in NCBI under BioProject ID PRJNA1536420, and via accession numbers listed in **Supplementary Table 1**.

## ACKNOWLEDGEMENTS

We gratefully acknowledge Drs. Breck Duerkop and Katrine Whiteson for providing the phages Hi3, Ben, and Bop. We acknowledge Dr. Shinjiro Ojima and Wakana Yamashita of the Japan National Institute of Infectious Diseases for their assistance in designing primers for cloning of anti-phage defense genes. We also acknowledge Dr. Emma Mills for assisting with bioinformatic analyses, and Dr. Madison Stellfox for providing subject matter expertise.

---

**Supplementary Figure 1.**
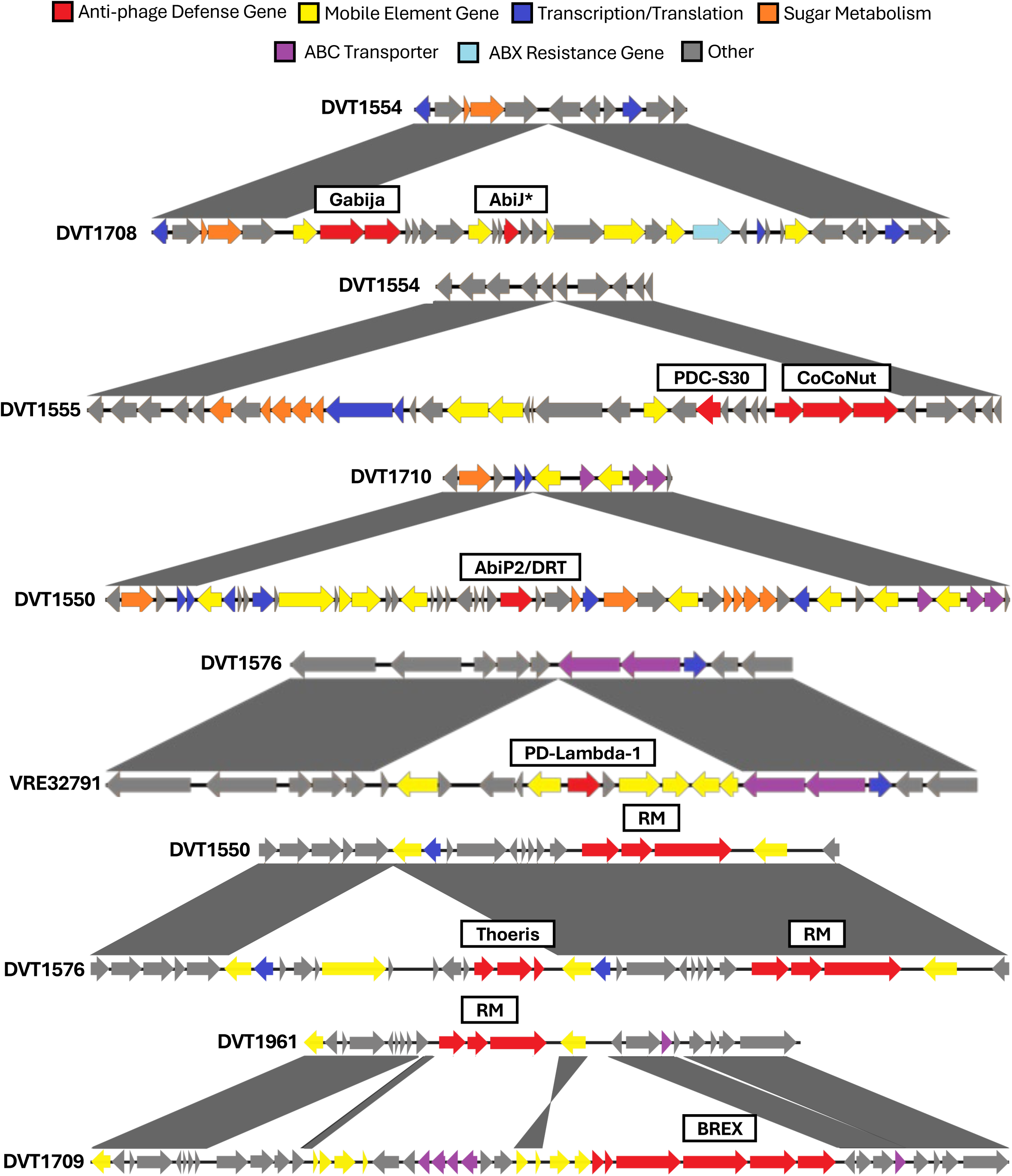
Visualization of defense islands in VREfm clinical isolate genomes. All islands shown are located on the bacterial chromosome, and genes are colored by functional annotation. Grey shading between sequences indicates homologous flanking regions. AbiJ* is an anti-phage defense system that was only identified when DefenseFinder was run with default search parameters.

**Supplementary Figure 2.**
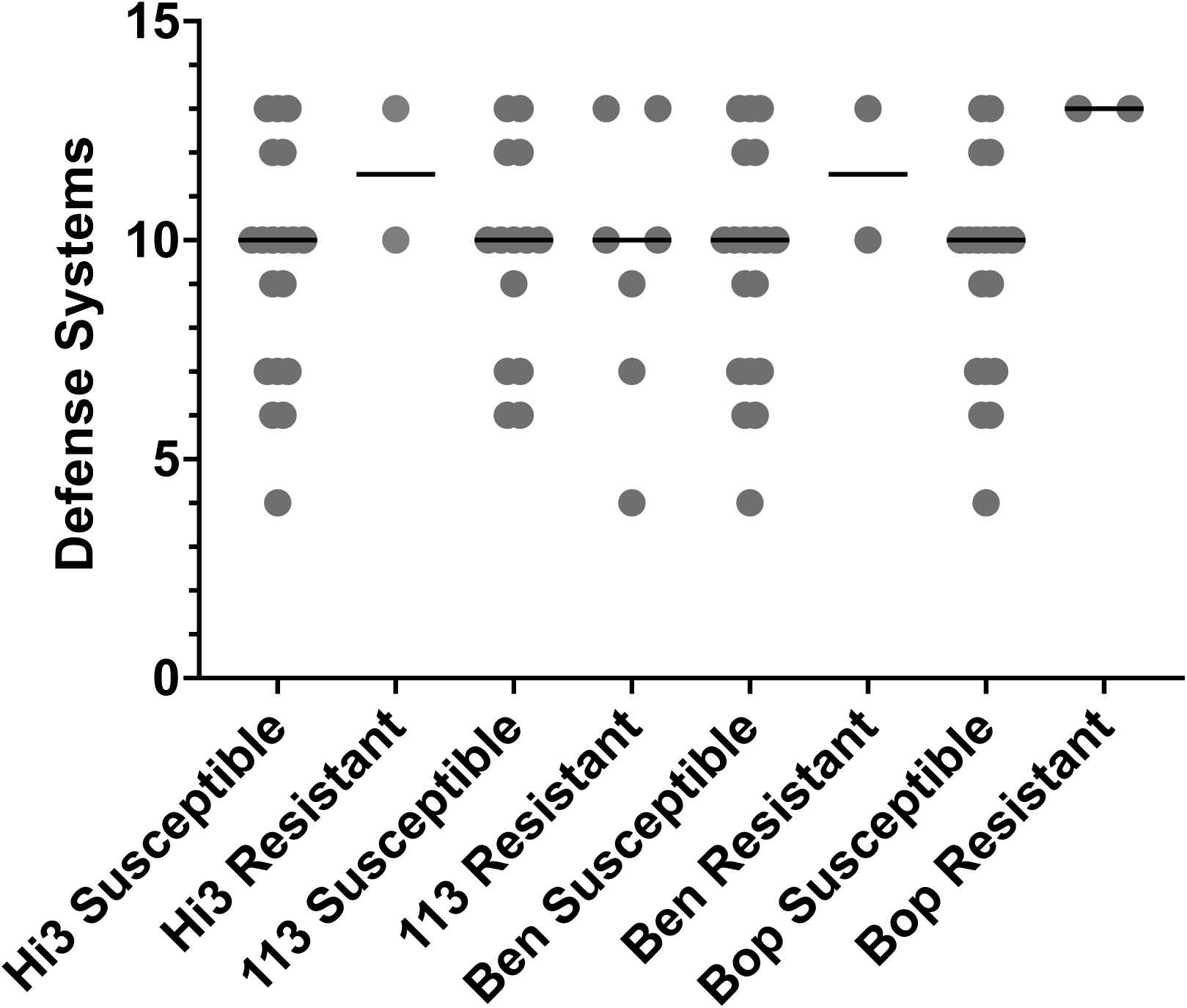
Number of intact anti-phage defense systems in genomes of phage-susceptible and phage-resistant VREfm isolates. Comparison of anti-phage defense system counts between isolates that were susceptible or resistant to each tested phage. Multiple copies of the same defense system were each counted individually.

**Supplementary Figure 3.**
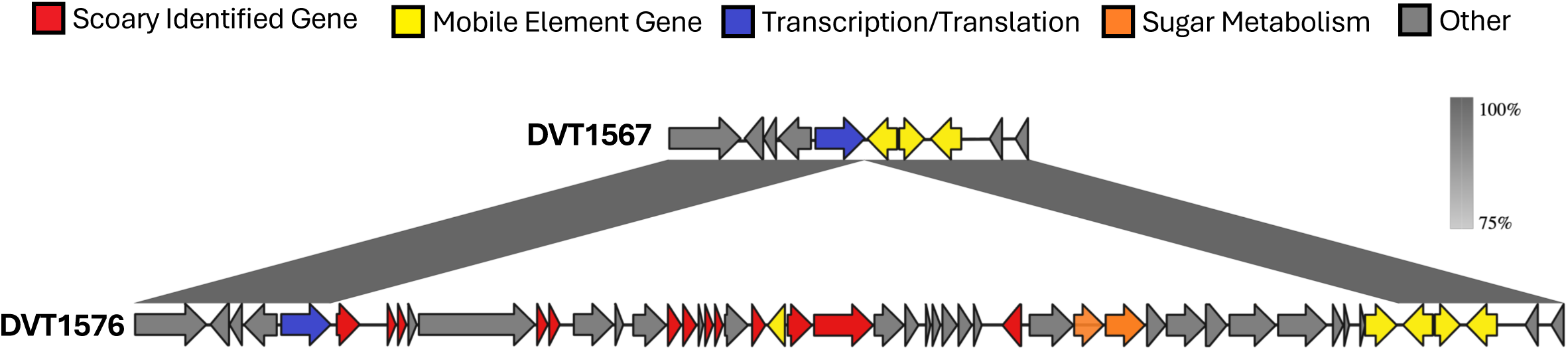
Putative defense island containing scoary-identified genes associated with phage killing. An isolate genome lacking the putative defense island is shown above an isolate encoding the island. Grey shading between sequences indicates homologous flanking regions and genes are colored by functional annotation.

## REFERENCES

1. Van Tyne D, Gilmore MS. 2014. Friend turned foe: evolution of enterococcal virulence and antibiotic resistance. Annu Rev Microbiol 68:337–56.

2. Arias CA, Murray BE. 2012. The rise of the Enterococcus: beyond vancomycin resistance. Nat Rev Microbiol 10:266–78.

3. Shay DK, Maloney SA, Montecalvo M, Banerjee S, Wormser GP, Arduino MJ, Bland LA, Jarvis WR. 1995. Epidemiology and mortality risk of vancomycin-resistant enterococcal bloodstream infections. J Infect Dis 172:993–1000.

4. CDC. 2019. Vancomycin-resistant Enterococci. cdcgov.

5. Fiore E, Van Tyne D, Gilmore MS. 2019. Pathogenicity of Enterococci. Microbiol Spectr 7.

6. Lebreton F, Manson AL, Saavedra JT, Straub TJ, Earl AM, Gilmore MS. 2017. Tracing the Enterococci from Paleozoic Origins to the Hospital. Cell 169:849–861 e13.

7. Edmond MB, Ober JF, Dawson JD, Weinbaum DL, Wenzel RP. 1996. Vancomycin-resistant enterococcal bacteremia: natural history and attributable mortality. Clin Infect Dis 23:1234–9.

8. Contreras GA, Munita JM, Simar S, Luterbach C, Dinh AQ, Rydell K, Sahasrabhojane PV, Rios R, Diaz L, Reyes K, Zervos M, Misikir HM, Sanchez-Petitto G, Liu C, Doi Y, Abbo LM, Shimose L, Seifert H, Gudiol C, Barberis F, Pedroza C, Aitken SL, Shelburne SA, van Duin D, Tran TT, Hanson BM, Arias CA. 2022. Contemporary Clinical and Molecular Epidemiology of Vancomycin-Resistant Enterococcal Bacteremia: A Prospective Multicenter Cohort Study (VENOUS I). Open Forum Infect Dis 9:ofab616.

9. Cheah AL, Spelman T, Liew D, Peel T, Howden BP, Spelman D, Grayson ML, Nation RL, Kong DC. 2013. Enterococcal bacteraemia: factors influencing mortality, length of stay and costs of hospitalization. Clin Microbiol Infect 19:E181–9.

10. Sangiorgio G, Calvo M, Migliorisi G, Campanile F, Stefani S. 2024. The Impact of Enterococcus spp. in the Immunocompromised Host: A Comprehensive Review. Pathogens 13.

11. Clokie MR, Millard AD, Letarov AV, Heaphy S. 2011. Phages in nature. Bacteriophage 1:31–45.

12. Johri AV, Johri P, Hoyle N, Pipia L, Nadareishvili L, Nizharadze D. 2021. Case Report: Chronic Bacterial Prostatitis Treated With Phage Therapy After Multiple Failed Antibiotic Treatments. Front Pharmacol 12:692614.

13. Paul K, Merabishvili M, Hazan R, Christner M, Herden U, Gelman D, Khalifa L, Yerushalmy O, Coppenhagen-Glazer S, Harbauer T, Schulz-Jurgensen S, Rohde H, Fischer L, Aslam S, Rohde C, Nir-Paz R, Pirnay JP, Singer D, Muntau AC. 2021. Bacteriophage Rescue Therapy of a Vancomycin-Resistant Enterococcus faecium Infection in a One-Year-Old Child following a Third Liver Transplantation. Viruses 13.

14. Stellfox ME, Fernandes C, Shields RK, Haidar G, Hughes Kramer K, Dembinski E, Mangalea MR, Arya G, Canfield GS, Duerkop BA, Van Tyne D. 2024. Bacteriophage and antibiotic combination therapy for recurrent Enterococcus faecium bacteremia. mBio 15:e0339623.

15. Aslam S, Lampley E, Wooten D, Karris M, Benson C, Strathdee S, Schooley RT. 2020. Lessons Learned From the First 10 Consecutive Cases of Intravenous Bacteriophage Therapy to Treat Multidrug-Resistant Bacterial Infections at a Single Center in the United States. Open Forum Infect Dis 7:ofaa389.

16. Djebara S, Maussen C, De Vos D, Merabishvili M, Damanet B, Pang KW, De Leenheer P, Strachinaru I, Soentjens P, Pirnay JP. 2019. Processing Phage Therapy Requests in a Brussels Military Hospital: Lessons Identified. Viruses 11.

17. Alrafaie AM, Pyrzanowska K, Smith EM, Partridge DG, Rafferty J, Mesnage S, Shepherd J, Stafford GP. 2024. A diverse set of Enterococcus-infecting phage provides insight into phage host-range determinants. Virus Res 347:199426.

18. Canfield GS, Chatterjee A, Espinosa J, Mangalea MR, Sheriff EK, Keidan M, McBride SW, McCollister BD, Hang HC, Duerkop BA. 2025. Correction for Canfield et al., “Lytic Bacteriophages Facilitate Antibiotic Sensitization of Enterococcus faecium”. Antimicrob Agents Chemother 69:e0059025.

19. Melo LDR, Ferreira R, Costa AR, Oliveira H, Azeredo J. 2021. Author Correction: Efficacy and safety assessment of two enterococci phages in an in vitro biofilm wound model. Sci Rep 11:11008.

20. Canfield GS, Duerkop BA. 2020. Molecular mechanisms of enterococcal-bacteriophage interactions and implications for human health. Curr Opin Microbiol 56:38–44.

21. Doron S, Melamed S, Ofir G, Leavitt A, Lopatina A, Keren M, Amitai G, Sorek R. 2018. Systematic discovery of antiphage defense systems in the microbial pangenome. Science 359.

22. Makarova KS, Wolf YI, Snir S, Koonin EV. 2011. Defense islands in bacterial and archaeal genomes and prediction of novel defense systems. J Bacteriol 193:6039–56.

23. Unterholzner SJ, Poppenberger B, Rozhon W. 2013. Toxin-antitoxin systems: Biology, identification, and application. Mob Genet Elements 3:e26219.

24. Perez Taboada V, Wu Y, Cassidy R, Medvedev KE, Loeff L, Nemudraia A, Nemudryi A. 2026. Bacterial Schlafen proteins mediate phage defence. Nat Microbiol 11:1037–1048.

25. Zhang T, Lyu Y, Beck CR, Iqbal N, Barbosa R, Ghanbarpour A, Laub MT. 2026. Bacterial immune activation via supramolecular assembly with phage triggers. Nature 651:1051–1059.

26. Vassallo CN, Doering CR, Littlehale ML, Teodoro GIC, Laub MT. 2024. Author Correction: A functional selection reveals previously undetected anti-phage defence systems in the E. coli pangenome. Nat Microbiol 9:2760–2761.

27. Garvey P, Rince A, Hill C, Fitzgerald GF. 1997. Identification of a recA homolog (recALP) on the conjugative lactococcal phage resistance plasmid pNP40: evidence of a role for chromosomally encoded recAL in abortive infection. Applied and Environmental Microbiology 63:1244–1251.

28. Chilambi GS, Nordstrom HR, Evans DR, Ferrolino JA, Hayden RT, Maron GM, Vo AN, Gilmore MS, Wolf J, Rosch JW, Van Tyne D. 2020. Evolution of vancomycin-resistant Enterococcus faecium during colonization and infection in immunocompromised pediatric patients. Proc Natl Acad Sci U S A 117:11703–11714.

29. Mills EG, Hewlett K, Smith AB, Griffith MP, Pless L, Sundermann AJ, Harrison LH, Zackular JP, Van Tyne D. 2025. Bacteriocin production facilitates nosocomial emergence of vancomycin-resistant Enterococcus faecium. Nat Microbiol 10:871–881.

30. Kunz Coyne AJ, Stamper K, Kebriaei R, Holger DJ, El Ghali A, Morrisette T, Biswas B, Wilson M, Deschenes MV, Canfield GS, Duerkop BA, Arias CA, Rybak MJ. 2022. Phage Cocktails with Daptomycin and Ampicillin Eradicates Biofilm-Embedded Multidrug-Resistant Enterococcus faecium with Preserved Phage Susceptibility. Antibiotics (Basel) 11.

31. Wandro S, Ghatbale P, Attai H, Hendrickson C, Samillano C, Suh J, Dunham SJB, Pride DT, Whiteson K. 2022. Phage Cocktails Constrain the Growth of Enterococcus. mSystems 7:e0001922.

32. Wick RR, Judd LM, Gorrie CL, Holt KE. 2017. Unicycler: Resolving bacterial genome assemblies from short and long sequencing reads. PLoS Comput Biol 13:e1005595.

33. Prjibelski A, Antipov D, Meleshko D, Lapidus A, Korobeynikov A. 2020. Using SPAdes De Novo Assembler. Curr Protoc Bioinformatics 70:e102.

34. Nishimura Y, Yoshida T, Kuronishi M, Uehara H, Ogata H, Goto S. 2017. ViPTree: the viral proteomic tree server. Bioinformatics 33:2379–2380.

35. Letunic I, Bork P. 2024. Interactive Tree of Life (iTOL) v6: recent updates to the phylogenetic tree display and annotation tool. Nucleic Acids Research 52:W78–W82.

36. Seemann T. mlst. https://github.com/tseemann/mlst. Accessed

37. Seemann T. 2014. Prokka: rapid prokaryotic genome annotation. Bioinformatics 30:2068–9.

38. Page AJ, Cummins CA, Hunt M, Wong VK, Reuter S, Holden MTG, Fookes M, Falush D, Keane JA, Parkhill J. 2015. Roary: rapid large-scale prokaryote pan genome analysis. Bioinformatics 31:3691–3693.

39. Stamatakis A. 2014. RAxML version 8: a tool for phylogenetic analysis and post-analysis of large phylogenies. Bioinformatics 30:1312–1313.

40. Stachurska X, Roszak M, Jablonska J, Mizielinska M, Nawrotek P. 2021. Double-Layer Agar (DLA) Modifications for the First Step of the Phage-Antibiotic Synergy (PAS) Identification. Antibiotics (Basel) 10.

41. Tesson F, Herve A, Mordret E, Touchon M, d’Humieres C, Cury J, Bernheim A. 2022. Systematic and quantitative view of the antiviral arsenal of prokaryotes. Nat Commun 13:2561.

42. Néron B, Denise R, Coluzzi C, Touchon M, Rocha EPC, Abby SS. 2023. MacSyFinder v2: Improved modelling and search engine to identify molecular systems in genomes. Peer Community Journal 3.

43. Tesson F, Planel R, Egorov AA, Georjon H, Vaysset H, Brancotte B, Néron B, Mordret E, Atkinson GC, Bernheim A, Cury J. 2024. A Comprehensive Resource for Exploring Antiphage Defense: DefenseFinder Webservice,Wiki and Databases. Peer Community Journal 4.

44. Mordret E, Herve A, Tesson F, Vaysset H, Clabby T, Loubat A, Shomar H, Planel R, Lavenir R, Cury J, Bernheim A. 2026. Protein and genomic language models uncover the unexplored diversity of bacterial immunity. Science 392:eadv8275.

45. Payne LJ, Meaden S, Mestre MR, Palmer C, Toro N, Fineran PC, Jackson SA. 2022. PADLOC: a web server for the identification of antiviral defence systems in microbial genomes. Nucleic Acids Res 50:W541–W550.

46. Payne LJ, Todeschini TC, Wu Y, Perry BJ, Ronson CW, Fineran PC, Nobrega FL, Jackson SA. 2021. Identification and classification of antiviral defence systems in bacteria and archaea with PADLOC reveals new system types. Nucleic Acids Res 49:10868–10878.

47. Sullivan MJ, Petty NK, Beatson SA. 2011. Easyfig: a genome comparison visualizer. Bioinformatics 27:1009–10.

48. Page AJ, Cummins CA, Hunt M, Wong VK, Reuter S, Holden MT, Fookes M, Falush D, Keane JA, Parkhill J. 2015. Roary: rapid large-scale prokaryote pan genome analysis. Bioinformatics 31:3691–3.

49. Wishart DS, Han S, Saha S, Oler E, Peters H, Grant JR, Stothard P, Gautam V. 2023. PHASTEST: faster than PHASTER, better than PHAST. Nucleic Acids Res 51:W443–W450.

50. Wang M, Liu G, Liu M, Tai C, Deng Z, Song J, Ou HY. 2024. ICEberg 3.0: functional categorization and analysis of the integrative and conjugative elements in bacteria. Nucleic Acids Res 52:D732–D737.

51. Bryan EM, Bae T, Kleerebezem M, Dunny GM. 2000. Improved vectors for nisin-controlled expression in gram-positive bacteria. Plasmid 44:183–90.

52. Shepard BD, Gilmore MS. 1995. Electroporation and efficient transformation of Enterococcus faecalis grown in high concentrations of glycine. Methods Mol Biol 47:217–26.

53. Brynildsrud O, Bohlin J, Scheffer L, Eldholm V. 2016. Rapid scoring of genes in microbial pan-genome-wide association studies with Scoary. Genome Biol 17:238.

